# Integration and analysis of CPTAC proteomics data in the context of cancer genomics in the cBioPortal

**DOI:** 10.1101/247718

**Authors:** Pamela Wu, Zachary J Heins, James T Muller, Adam A Abeshouse, Yichao Sun, Nikolaus Schultz, David Fenyö, Jianjiong Gao

## Abstract

The Clinical Proteomic Tumor Analysis Consortium (CPTAC) has produced extensive mass spectrometry based proteomics data for selected breast, colon and ovarian tumors from The Cancer Genome Atlas (TCGA). We have incorporated the CPTAC proteomics data into the cBioPotal to support easy exploration and integrative analysis of these proteomic datasets in the context of the clinical and genomics data from the same tumors. cBioPortal is an open source platform for exploring, visualizing, and analyzing multi-dimensional cancer genomics and clinical data. The public instance of the cBioPortal (http://cbioportal.org/) hosts more than 100 cancer genomics studies including all of the data from TCGA. Its biologist-friendly interface provides many rich analysis features, including a graphical summary of gene-level data across multiple platforms, correlation analysis between genes or other data types, survival analysis, and network visualization. Here, we present the integration of the CPTAC mass spectrometry based proteomics data into the cBioPortal, consisting of 77 breast, 95 colorectal, and 174 ovarian tumors that already have been profiled by TCGA for mutations, copy number alterations, gene expression, and DNA methylation. As a result, the CPTAC data can now be easily explored and analyzed in the cBioPortal in the context of clinical and genomics data. By integrating CPTAC data into cBioPortal, limitations of TCGA proteomics array data can be overcome while also providing a user-friendly web interface, a web API and an R client to query the mass spectrometry data together with genomic, epigenomic, and clinical data.

## Introduction

In the last decade, The Cancer Genome Atlas (TCGA) consortium has generated multi-platform cancer genomics data, including somatic mutations, copy number alterations, gene expression, and DNA methylation, in more than 20 cancer types (1). TCGA also generated some proteomics data using the Reverse Phase Protein Array (RPPA) platform, measuring protein levels in tumors for about 150 proteins and 50 phosphoproteins (2). However, the RPPA technology is limited by the availability and binding efficiency of available antibodies for protein and post-translational modification detection. The Clinical Proteomic Tumor Analysis Consortium (CPTAC) consortium is aimed at characterizing the protein inventory in tumors by leveraging the latest developments in mass spectrometry-based discovery proteomics. The initial CPTAC projects have generated extensive mass spectrometry-based proteomics data on TCGA tumors of breast cancer (proteome and phosphoproteome) (3, 4), ovarian cancer (proteome, phosphoproteome, and glycoproteome) (5, 6), and colorectal cancer (proteome only, with matching normal samples) (7, 8, 9). Currently, the consortium is extending the proteomics analysis to genomically characterized tumor samples for other cancer types.

By profiling the same cancer patients already profiled by TCGA, CPTAC results provide a unique opportunity for performing integrative analysis of cancer genomics and proteomics data, which can link proteomics to genotypes and potentially phenotypes in cancer. There is a growing demand for exploratory analysis tools that integrate cancer genomics, proteomics and clinical data and provide easy access to the multi-dimensional datasets, and efforts such as TCPA and LinkedOmics have started to fill this gap (10, 11). The cBioPortal for Cancer Genomics has been one of the leading resources for analyzing cancer genomics data, including all TCGA projects and many data sets curated from the literature (12, 13). cBioPortal features a biologist-friendly interface, biology-aware visualization, and integrative analysis features, making it one of the most popular resources in the community of cancer genomics researchers, especially biologists without bioinformatics skills.

We have updated cBioPortal to integrate the CPTAC mass spectrometry based proteomics data, making the high quality proteomics data easily accessible for visualization and analysis in the context of cancer genomics. First, we developed a data transformation pipeline for converting mass spectrometry results produced by CPTAC members into a data format that is compatible with the cBioPortal data pipeline. This data transformation pipeline is also applicable for uploading non-CPTAC mass spectrometry results into cBioPortal, as they become available. In order to better support integrative analysis of genomics and proteomics data, we have made various adjustments to the cBioPortal interface, including protein expression heatmap visualization in OncoPrint (graphic summary of gene-level data and clinical attributes), correlation analysis, and differential analysis. By integrating CPTAC data into cBioPortal, we aim to increase the accessibility of mass spectrometry-based proteomics data to cancer researchers in the context of cancer genomics and provide researchers with an intuitive interface to explore and analyze interactions between genomics and proteomics in cancer.

## Experimental Procedures and Results

### Introduction to the cBioPortal for cancer genomics interface

The public instance of the cBioPortal hosts more than 100 studies, including all TCGA projects and published studies from the literature. A key step in cBioPortal that enables integration of various datasets is gene-level mapping of the data. For example, by mapping mutations, copy number, and gene expression onto a gene, the visualization and analysis features of the cBioPortal supports studying different types of gene alterations in tumors simultaneously, including, among others:

- **Gene-oriented query**: A simple web form that allow users to query alterations to genes of interest in individuals or across cancer studies. Samples were classified into the *altered group* (if there is an alteration in any of the query genes in the sample) or the *unaltered group* for downstream analysis.
- **OncoPrint**: A graphical summary of genomic alterations across samples in multiple genes, represented by different glyphs and color coding. This graphical representation gives an overview of genomic alterations to the genes of interest in a selected cohort. Clinical attributes can also be visualized together with the genomics data. To better support visualization of proteomics data, we have added heatmap visualization into OncoPrint.
- **Correlation analysis**: The Plots tab can be used to perform correlation analysis between genomics data (copy number alteration and gene expression), epigenomics data (DNA methylation), proteomics data (RPPA, and now mass spectrometry based data), and clinical attributes for a gene or between two genes. Mutation data is overlaid onto the correlation plots.
- **Co-expression analysis**: For each queried gene, Pearson’s and Spearman’s correlation coefficients are calculated against all other genes in a selected gene expression profile.
- **Enrichments analysis**: After classifying samples into altered and unaltered groups based on the query, this analysis identifies mutations or copy number alterations that are enriched in either group (by a Fisher’s exact test). It also identifies genes and proteins that are over/under-expressed in a group (by a two-sided, two-sample Student’s t-test).
- **Survival analysis**: Kaplan-Meier estimators and plots for overall survival and disease-free or progression-free survival are generated to study the survival difference between altered and unaltered groups.
- **Mutual exclusivity analysis**: Fisher’s exact test to analyze whether alterations are significantly mutually exclusive or co-occurring between every pair of query genes.
- **Patient view page**: Summary and visualization of clinical attributes and genomic alterations in a tumor or a patient.
- **Study view page**: Summary, visualization, and interactive exploration of clinical attributes and genomic alterations in a cohort of tumors.

### Transforming CPTAC data for the cBioPortal database

Processed CPTAC mass spectrometry data was downloaded from the CPTAC Data Portal (https://cptac-data-portal.georgetown.edu/). The downloaded data had been processed using the Common Data Analysis Pipeline (CDAP) (14), which standardizes the treatment of data across the consortium. CDAP software is flexible to accommodate different types of mass spectrometry data. Peptides are identified using MS-GF+ (15), and the protein FASTA file used is a concatenation of RefSeq *H. sapiens* build 37 with the sequence for S. scrofa trypsinogen added. The MS-GF+ phosphopeptide assignments are fed into PhosphoRS to obtain phosphosite localizations. The results are made into both peptide-level and protein-level tab-separated (TSV) files consisting of iTRAQ ratios for breast and ovarian sample and precursor area intensity measurements for the colon samples.

We mapped the proteomics data onto genes because the query interface and analysis features of cBioPortal are gene-centric, so protein levels were mapped directly to their corresponding genes by mapping RefSeq protein IDs to Hugo gene symbols. For PTM data, w
e focused on the phosphoproteomics data. For phosphoprotein levels (currently, the only PTM added to cBioPortal), we make special symbols for them with the following pattern:

> `<HUGO gene symbol>_P{S,T,Y}<AA modified>`

For example, EIF4EBP1 phosphorylated at the serine located at position 65 is denoted as EIF4EBP1_PS65. In order to easily access phosphoproteins while querying, each phosphoprotein in the database has as an alias

> `PHOSPHO <HUGO gene symbol>`

which, when queried, brings up a list of phosphoproteins sharing that alias. Users can choose the phosphoproteins that they want to see displayed. Future PTMs will be associated with similar aliases as they are added.

In order for the data to be used by cBioPortal, the distribution of intensities per sample must be reasonably close to a normal distribution, since the interface relies on Z-scores for much of its computations and displays. For both the iTRAQ and the label-free quantitation data, the log-transformed intensities are approximately normally distributed, and therefore their values were kept as intensities. In addition, the global proteomics and posttranslational modification (PTM) data were formatted as required by the cBioPortal import pipeline.

Additionally, we also implemented an option of converting MS intensity values into estimates of protein copy number per cell using the `-proteomic-ruler` parameter. This function uses the methods outlined in Wiśniewski et al. 2014 (16) to transform protein intensities into copy number values by taking advantage of the observation that DNA and histone mass are fairly constant per cell and that the ratio between protein and histone mass is proportional to the ratio between protein and histone mass spectrometry signals. The equation they have outlined is as follows:

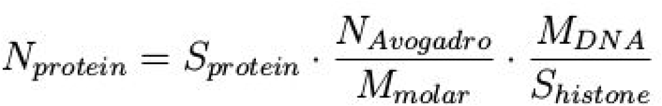

N stands for number (Avogadro’s number, 6.022×10^23^, and the measured protein’s copy number), S stands for mass spectrometry signal value (of histones and the measured protein), and M stands for mass (of DNA per cell which is estimated to be 6.5 pg (16) and the molar mass of the measured protein). cBioPortal currently only hosts the raw, unprocessed intensities, but this feature for the processing pipeline has been added to increase the interpretability of intensity values of mass spectrometry data.

The classes created for the processing of CDAP and MaxQuant proteome and PTM file formats can be found on GitHub (see Supplementary Information 5), which also includes a usage tutorial that leads users through the API. For MaxQuant proteome data, which is generally contained in the “proteinGroups.txt” file, intensity data is filtered for q-values under 0.05. MaxQuant PTM data is filtered for a localization probability of over 0.75.

### Use case 1: The cBioPortal web interface facilitates exploration of regulatory patterns of mRNA and protein expression data of cancer patients

The CPTAC proteomics data can now be interactively analyzed and visualized in the context of TCGA genomics and clinical data in the cBioPortal. We have added the new proteomics data in the query interface as a new genetic profile (currently for TCGA breast, colorectal, and ovarian provisional studies), which allows for querying the protein and phosphoprotein levels together with gene-level genomics data (mutations, copy number alterations, and gene expression). After submitting a query, the proteomics data has been incorporated in downstream visualization and analysis features.

One new feature we added was to support the generation of a heatmap of expression data in the OncoPrint tab. Upon query submission, the user now has an option to append heatmaps of gene and protein expression to the OncoPrint summarizing genomic alterations of the query genes and samples. The new heatmap feature allows researchers to identify expression patterns at a glance. For example, it is possible to view the protein levels of a gene and its identified phosphosites and cluster the data along both dimensions (Figure 1). These new features can be combined with previously released features to gain an integrative overview of genomic and clinical attributes of the patient samples, which can be used to easily identify data trends or individual patient samples of interest. For example, in Figure 1, it can be observed that ERBB2 copy number amplification, mRNA expression, and protein and phosphosite level detection are all very closely correlated with whether or not ERBB2 receptors are detected on the surface of tumors, as indicated by the HER2-IHC score. Also, TCGA-A2-A0T6 notably has an unusual mutation spectrum that is enriched for the thiamine to guanine transversion. The patient seems to have a lower than average diagnosis age and overall survival, and the patient has been classified as having breast cancer lobular carcinoma. Interestingly, this tumor sample has three oncogenic ERBB2 mutations (shown when hovering over the sample), and its ERBB2 is expressed above average in mRNA but below average in protein and all available phosphosites.

**Figure 1.**
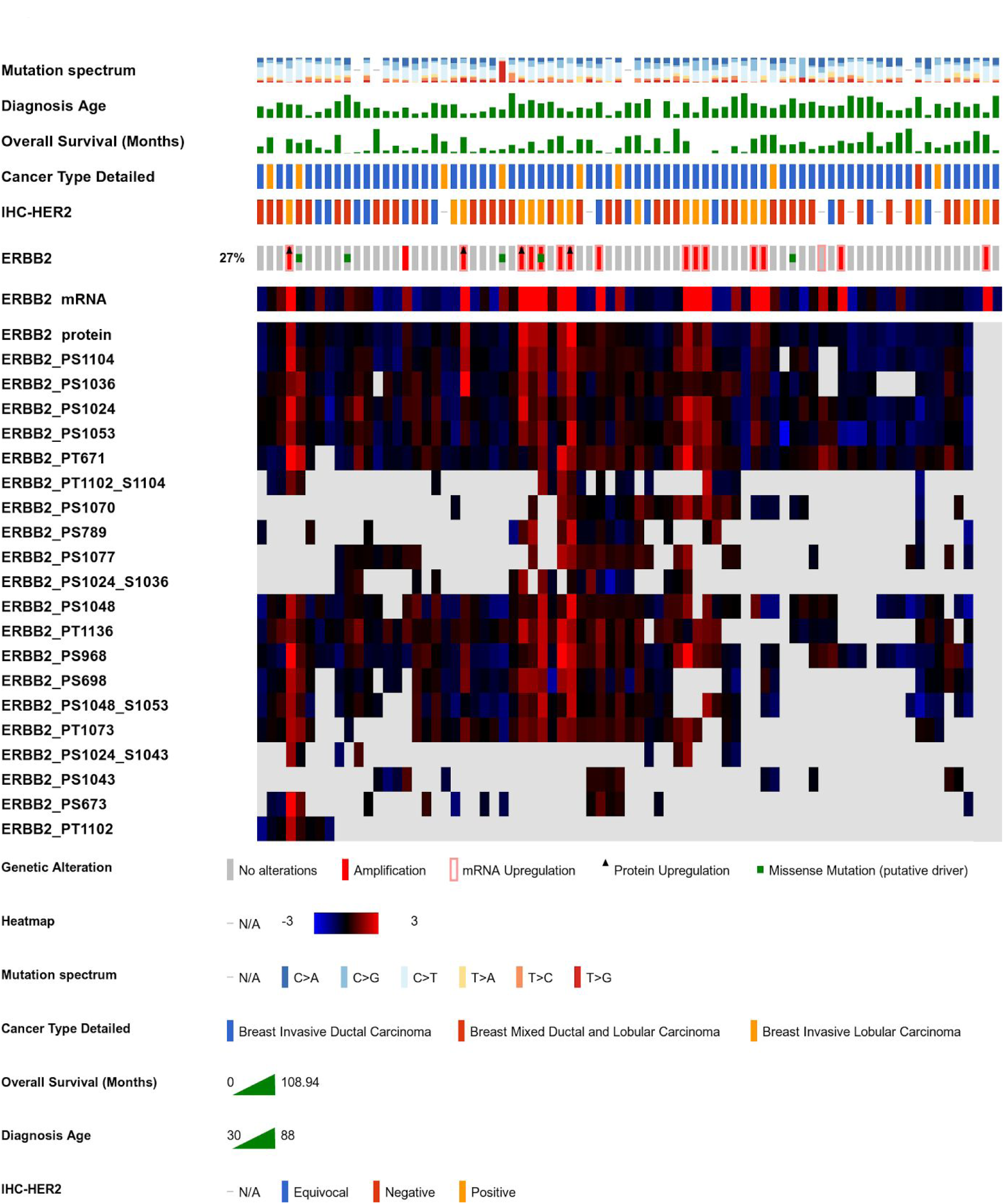
Oncoprint of ERBB2 alterations including heatmap of its protein levels and the levels of ERBB2 phosphosites. Samples consisted of the 77 TCGA breast cancer patient samples containing genomic alterations, RNAseq RSEM data, and mass spectrometry data for ERBB2 along with the mass spectrometry data for all available phosphosites for ERBB2 containing any non-NaN data across all samples. Clinical information for this example includes the mutation spectrum, which is a segmented bar plot of the fraction that each transition or transversion occurs within the sample (though not necessarily for the gene set specified); diagnosis age; overall survival, which is the survival in months from the time of diagnosis; the detailed cancer type, which in this case indicates if the breast cancer was lobular or ductal; and the HER2 (ERBB2) IHC score, which indicates whether or not immunohistochemistry (IHC) staining was able to detect ERBB2 receptors on the surface of the tumors (negative corresponds to a score of 0-1, equivocal corresponds to a score of 2, and 3+ is positive for ERBB2).

Other tabs can also allow users to explore potential correlations in depth. For example, the Plots tab can be used to select the DNA copy number, methylation, gene expression, or protein levels from the list of queried genes and post-translational modifications, which allows the user to browse the Pearson and Spearman correlations between the protein levels of ERBB2 and its phosphosites. By modifying the query to select other cancer types, it is also possible to see how these correlations shift across cancers (Figure S1).

To illustrate the value of data exploration using well-established examples, we centered on ERBB2, which is a known oncogene in breast and ovarian cancer. The gene sequences of ERBB2 and GRB7 lie on the same amplicon on chromosome 17q12-21 (17). It has been shown that GRB7 can act as an ERBB2-dependent oncogene by enhancing ERBB2 phosphorylation (18). Using the cBioPortal Plots tab, users can see a perfect co-occurrence of copy number categorizations between ERBB2 and GRB7. This persists as a high correlation between copy number and mRNA levels and mRNA and protein levels. ERBB4, on the other hand, is in the same receptor family as ERBB2, and it is not located on the same amplicon so copy number changes are not correlated. Also mRNA levels of of ERBB2 and ERBB4 are not correlated but protein levels are correlated consistent with ERBB2 and ERBB4 forming a heteromeric complex (Figure S2) (19).

### Use case 2: Exploring cBioPortal data through its web API using the R package cgdsr to access the full database now loaded with mass spectrometry data

While the cBioPortal web interface offers many pre-built analysis tools for exploratory analysis, users may want to perform further custom and automated analysis of the data that cannot be done on the interface. The cBioPortal public database has integrated all the different genomic profiles from TCGA, CPTAC, and other sources together into a MySQL database, and made the data available programmatically via a web application program interface (API). The R package cgdsr accesses the all the genomics and clinical data by connecting to the web API. An example R script to access cBioPortal data including the proteomics data is provided here (see Supplementary Information 6). In this example, we looked at how well PAM50-based subtypes are conserved when using mass spectrometry-based proteomics values for clustering. Using the “`ward. D2`” setting for the `histfun` parameter of `heatmap .2` in R, we performed hierarchical clustering on the 77 breast cancer samples containing mass spectrometry data and color-coded the samples by their PAM50 subtype classification (see Figure S3). A more rigorous version of this analysis has been performed on the same data (4), but this example shows the streamlining of data retrieval and analysis that is achieved by wrapping the CPTAC data with the cBioPortal API (see Supplementary Script S1).

### Use case 3: Loading unpublished or institution-specific proteomics data onto a local private cBioPortal instance for integrative analysis

Over 30 institutions now have a private instance of cBioPortal hosting institution-specific research and/or clinical data, which allows researchers to perform cBioPortal-enabled exploratory analysis on private institutional data. An import pipeline now exists to convert mass spectrometry data processed by CDAP into a format for effective loading into the cBioPortal database. All features and functionalities available on the public interface is available for private institutional data which can be supplemented with the public data for richer exploratory analysis. Instructions for the deployment of a private instance can be found in the documentation (see Supplementary Information 7).

## Discussions and Conclusion

Being able to easily integrate multiple omics domains in order to synthesize them into actionable insights is a modern challenge created by the proliferation of high-throughput sequencing and spectrometry techniques. To address this challenge, the cBioPortal has been developed to provide a public and user-friendly interface for performing interactive and integrative data exploration, analysis and visualization. To incorporate the mass spectrometry data from CPTAC, we have updated the cBioPortal data pipeline and the public web interface. The data process pipeline not only enables integration of current and future CPTAC data, but also allows private mass spectrometry data to be loaded into databases of institutional instances of the cBioPortal. We also updated the web interface and web API of cBioPortal to support easy access and better visualization and analysis of the mass spectrometry data in the context of cancer genomics and clinical datasets.

The inclusion of mass spectrometry data from CPTAC in the cBioPortal is a significant step towards better understanding of the whole molecular profile of various cancer types. In TCGA projects, RPPA was adopted as the proteomics platform to profile key proteins and has generated a valuable resource for study cancer proteomics and its association with cancer genomics. However, due to the availability of antibodies, the coverage of RPPA for measuring the proteome (both total proteins and PTMs) is limited, while mass spectrometry can profile a large portion of the proteome and many different types of PTMs without the initial data collection biased by the availability of detection agents (for example, see comparison in Figure S4). At the moment, the number of tumors analyzed mass spectrometry is still considerably smaller than those analyzed by RPPA (see Table 1), but with the continuous efforts such as CPTAC, we expect to see a large increase of the number of tumors in more cancer types profiled by mass spectrometry. All in all, integrating mass spectrometry data with genomics data provides a new opportunity to uncover novel associations between genome, proteome and phenotype in cancer.

**Table 1:** Cancer studies in cBioPortal currently updated with the new mass spectrometry data. Query Menu Location refers to the location in the tree structure that organizes the cancer studies on the homepage of cBioPortal where the initial query parameters are set.

| Tissue | RPPA Samples | MS Samples | Query Menu Location |
| --- | --- | --- | --- |
| Breast | 892 | 77 | Breast → Invasive Breast Carcinoma → Breast Invasive Carcinoma (TCGA, Provisional) |
| Colorectal | 357 | 95 | Bowel → Colorectal Adenocarcinoma → Colorectal Adenocarcinoma (TCGA, Provisional) |
| Ovarian | 498 | 174 | Ovary → Ovarian Epithelial Tumor → Serous Ovarian Cancer → High-Grade Serous Ovarian Cancer → Ovarian Serous Cystadenocarcinoma (TCGA, Provisional) |

## Acknowledgements

PW was funded by the Google Summer of Code program and program development funds from New York University School of Medicine. DF was funded by National Cancer Institute (NCI) CPTAC award U24 CA210972 and by contract 13XS068 from Leidos Biomedical Research. JG
 and NS were funded by the Marie-Josée and Henry R. Kravis Center for Molecular Oncology, a National Cancer Institute Cancer Center Core Grant (P30-CA008748), and the Robertson Foundation (NS).

## Supplementary Information

The CPTAC Data Portal hosts all of the data produced by the consortium (https://cptac-data-portal.georgetown.edu/cptacPublic/). The data can also be obtained using HTTP (https://cptc-xfer.uis.georgetown.edu/publicData/). The public cBioPortal interface can be found on the main site (http://www.cbioportal.org/). The open source code base is hosted on GitHub under an AGPL-3.0 license (https://github.com/cBioPortal/cbioportal). Scripts for the production of import files for mass spectrometry data is also hosted on GitHub(https://github.com/cBioPortal/CPTAC-proteomics-pipeline) under an MIT license.

**Script S1.**
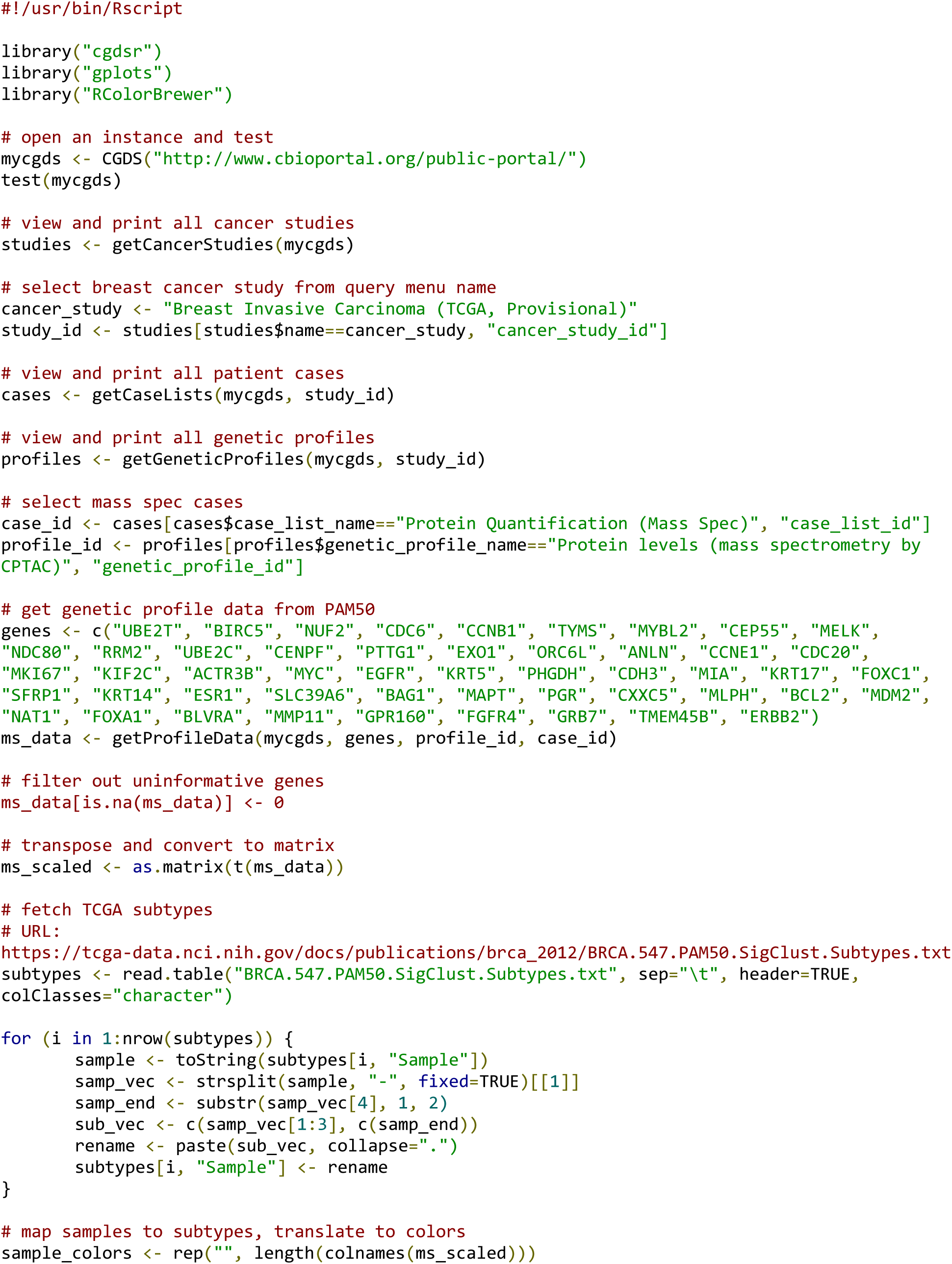
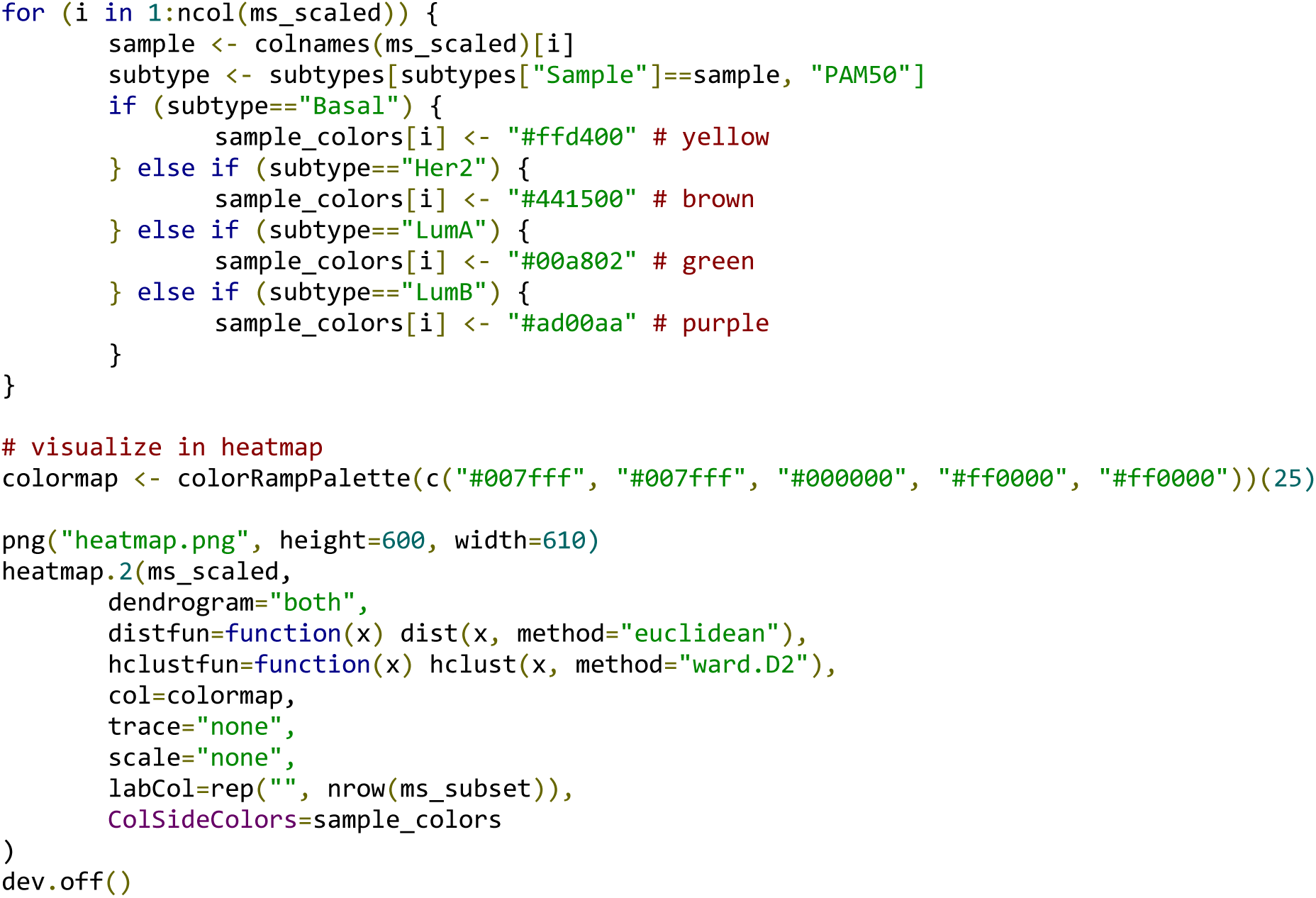

The script file used for generating the heatmap shown in Figure S3 is as follows:

**Figure S1.**
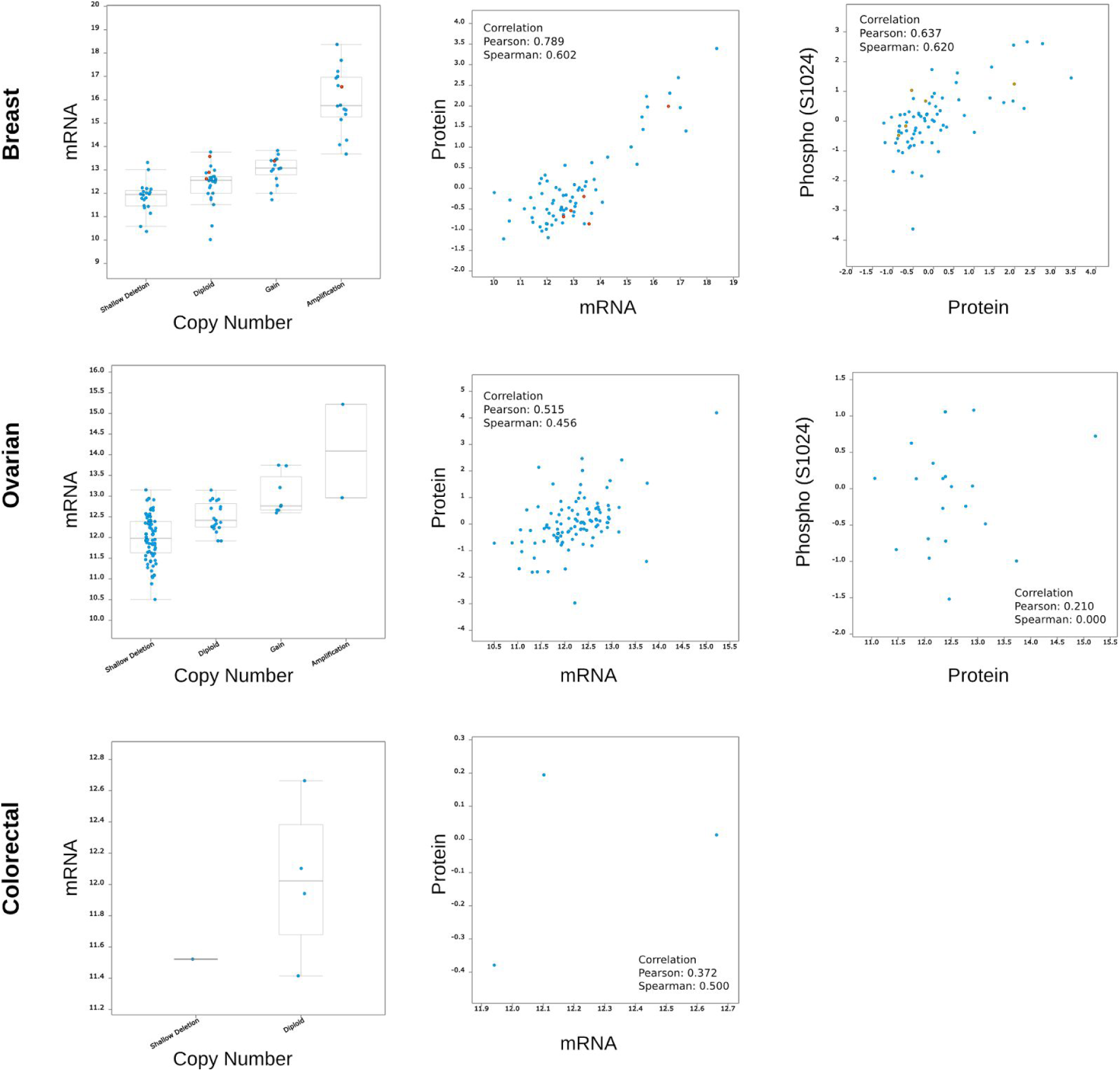
Correlation plots for co-occurrences and correlations between ERBB2 copy number, mRNA, and protein levels in breast, ovarian, and colorectal cancers. Categories of copy number types include deep deletions, shallow deletions, diploid, gain, and amplification. Both the Pearson and Spearman correlations are reported. This data incorporates the mass spectrometry data generated from TCGA samples by the CPTAC consortium. No phosphosite information is available for colorectal cancer from this dataset.

**Figure S2.**
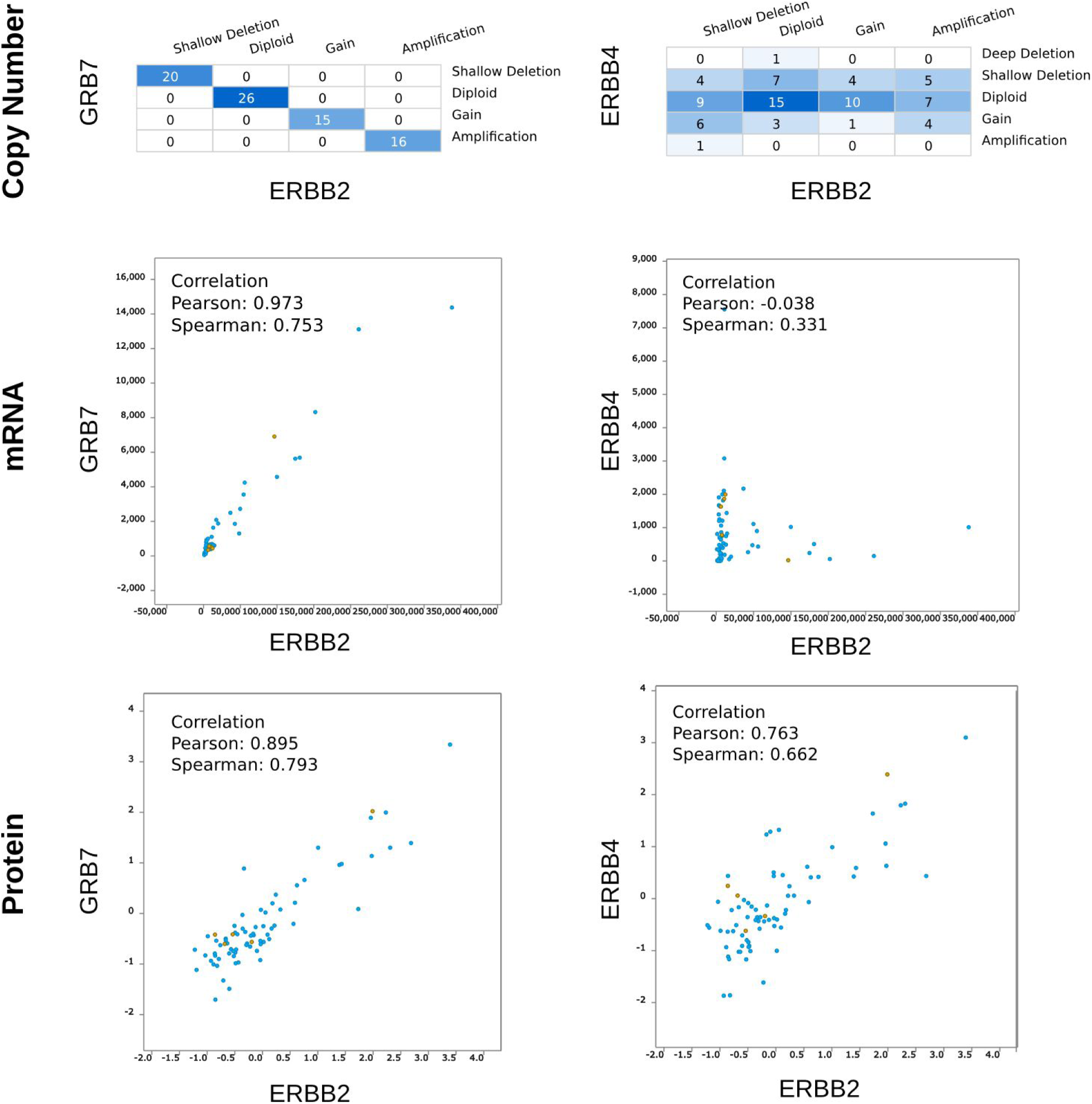
Correlation plots for co-occurrences and correlations between copy number, mRNA, and protein levels for ERBB2 vs. GRB7 and ERBB2 vs. ERBB4. Copy number plots are shown as co-occurrences and all other genomic profiles are shown as correlations.

**Figure S3.**
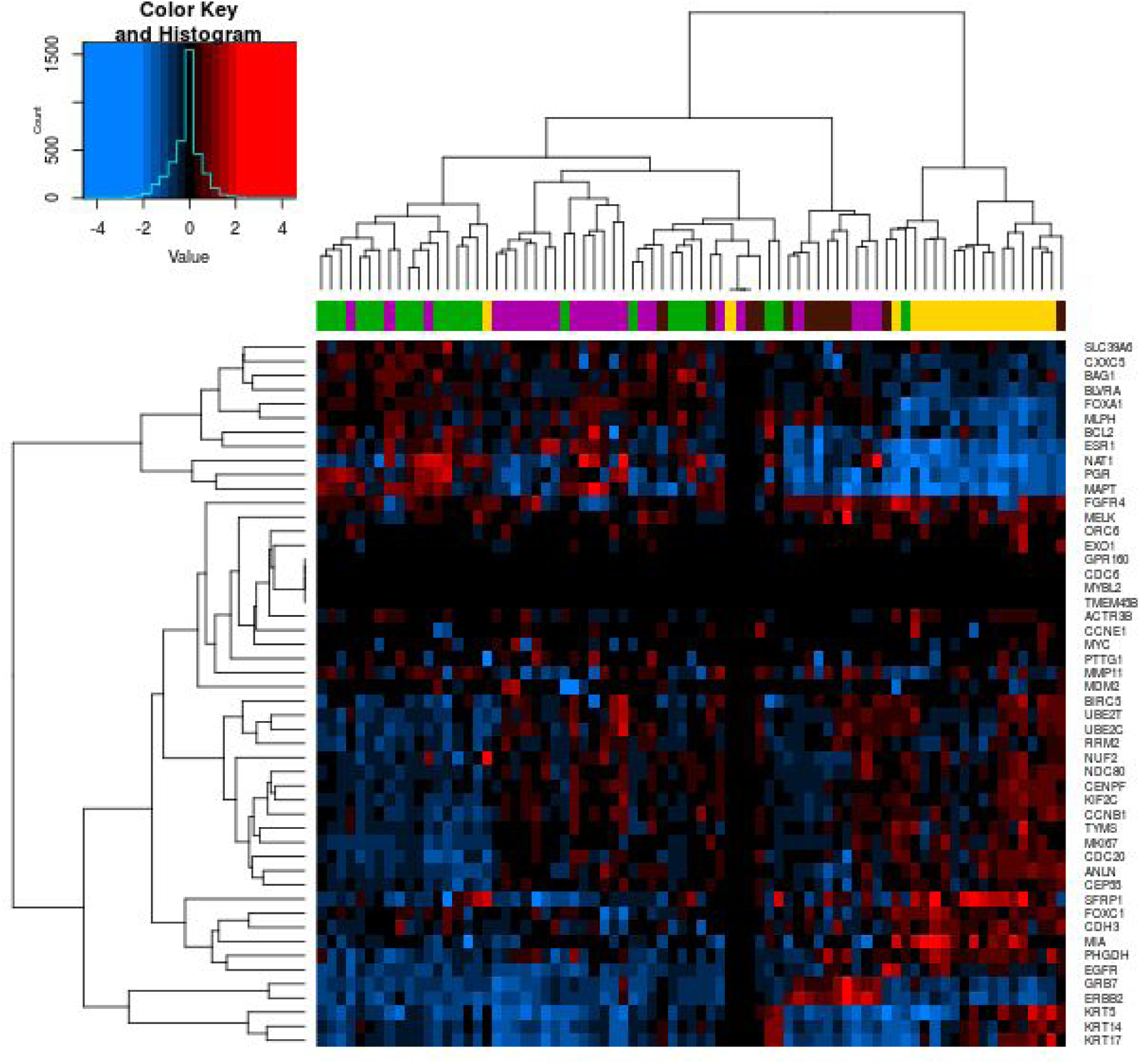
A heatmap of 77 breast cancer TCGA patient samples containing CPTAC mass spectrometry proteomics data, clustered by PAM50 genes. Data values are log2-transformed iTRAQ quantitation values for each gene and sample. Samples were color-coded by their original PAM50 subtype classification: Basal (yellow), Her2 (brown), Luminal A (green), and Luminal B (purple).

**Figure S4.**
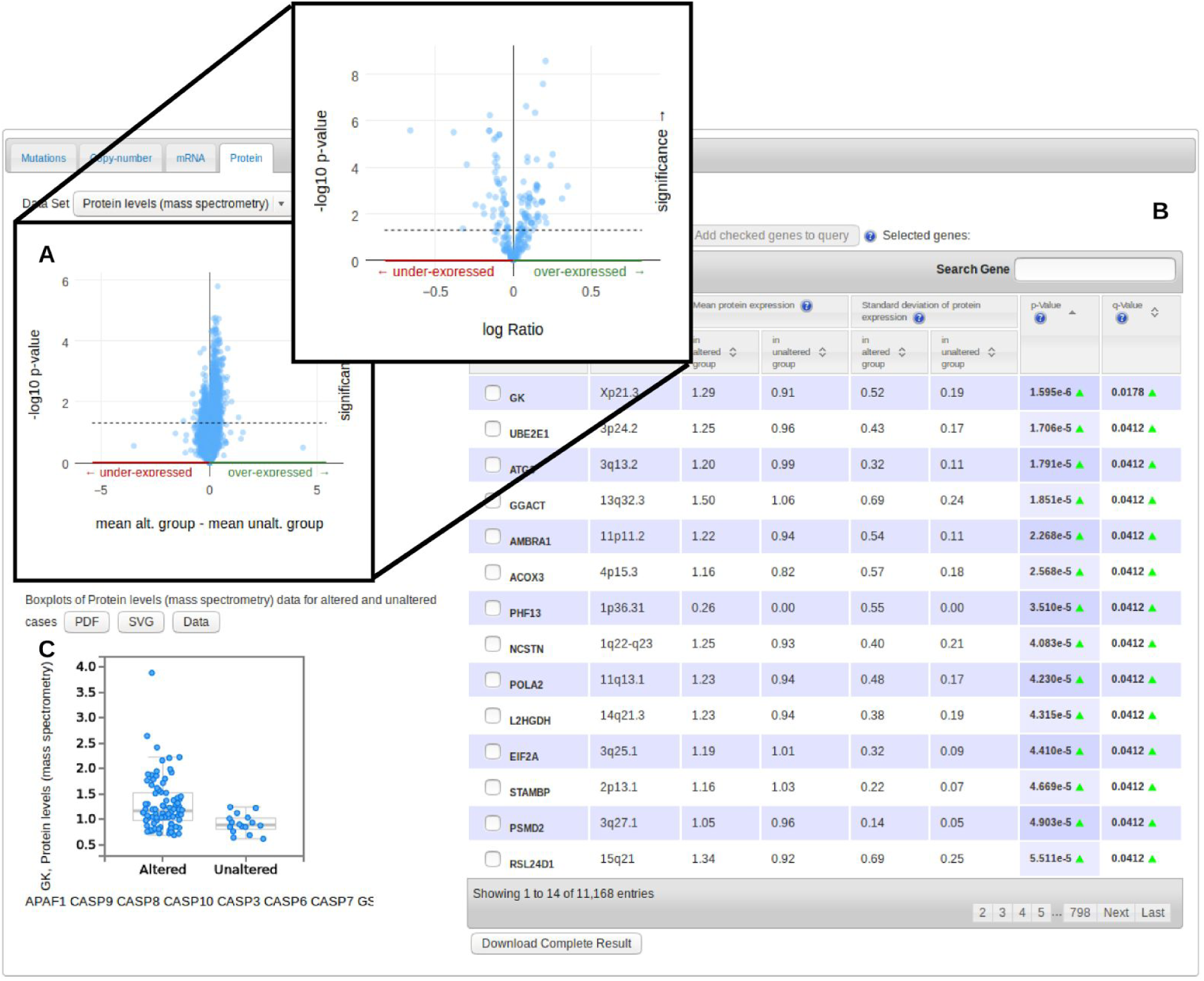
The Enrichment tab for protein expression from mass spectrometry data from 77 patients in the Breast Cancer Provisional TCGA study. a) A volcano plot of the protein expression levels from all genes with mass spectrometry data available. The pop-out shows the volcano plot for the RPPA data for the same query. b) A table of non-queried genes that show significant expression differences between altered and unaltered patients. c) A boxplot of the protein expression values between altered and unaltered patients for the gene GK, which was the top scoring gene by FDR-adjusted q-value.

